# Squid primary cell culture as a model system and experimental tool

**DOI:** 10.1101/2025.01.12.632648

**Authors:** Yoonji Kim, Hailey M. Tanner, Joshua J. C. Rosenthal, Clifford P. Brangwynne

## Abstract

Squid primary cells from various tissues and ages are isolated, maintained in culture, and express exogenous genes. This protocol opens up numerous opportunities in molecular biology, neuroscience, and marine biology, enabling molecular and cellular-level investigations into processes specific to squids, including their complex behaviors, rapid color change mechanisms, RNA editing capabilities, with broader implications in basic cell physiology. We also describe procedures that harness life-stage tractability in *Euprymna berryi* squids towards future studies of aging in this model marine organism.

**Highlights:**

- Squid primary cells are isolated from optic lobes, gills, eyes, and the skin
- Cells from these tissues can be dissociated from any life stage of the squid
- Isolated cells can express exogenous genes through mRNA transfections and are subject to live and/or fixed cell imaging
- Specific details for optimized media conditions, trypsinization, plating, and passaging are included

## Before you begin

Coleoid cephalopods–namely, the octopus, squid, and cuttlefish–are the most behaviorally sophisticated invertebrates with highly developed sensory and nervous systems and camera-type eyes. They display iridescence on their skin and have specialized chromatophores to initiate camouflage with their immediate surroundings. In addition to these unique adaptations and processes, including their complex cognitive behaviors and rapid color change mechanisms on the organismal level, cephalopods can edit their genetic information through A-to-I RNA editing and have made important contributions to the fields of neuroscience and molecular biology, with studies of the giant squid axon revealing that information travels electrically between neurons^1^ and identifying and isolating kinesin from the giant axons and optic lobes^2^.

The *Euprymna berryi (E. berryi)* bobtail hummingbird squid is an emerging new model organism amenable to genetic manipulation^3,4^. Bobtail squids have primarily been studied for their unique symbiotic relationships with bioluminescent bacteria *Vibrio fischeri*^5^ and likely use extensive A-to-I RNA editing to sense and adapt to their environment, similar to other cephalopod species^6^. *E. berryi* squids are ideal to study in the laboratory setting due to their small size (∼2 inches long), short time to reach sexual maturity, relatively short lifespan of 6-8 months and the ability to tolerate crowding. Another feature of these squids is the ability to monitor stereotyped life-stage changes over experimentally tractable life timescales (Figure 1A). Researchers are able to breed these squids, and due to their prolific and predictable reproductive cycles, they are a promising model organism to study aging. Additionally, unlike how octopuses senesce and die post brooding eggs^7^, female *E. berryi* squids do not and will continue to live and lay eggs until reaching an old age of 6-8 months.

**Figure 1:**
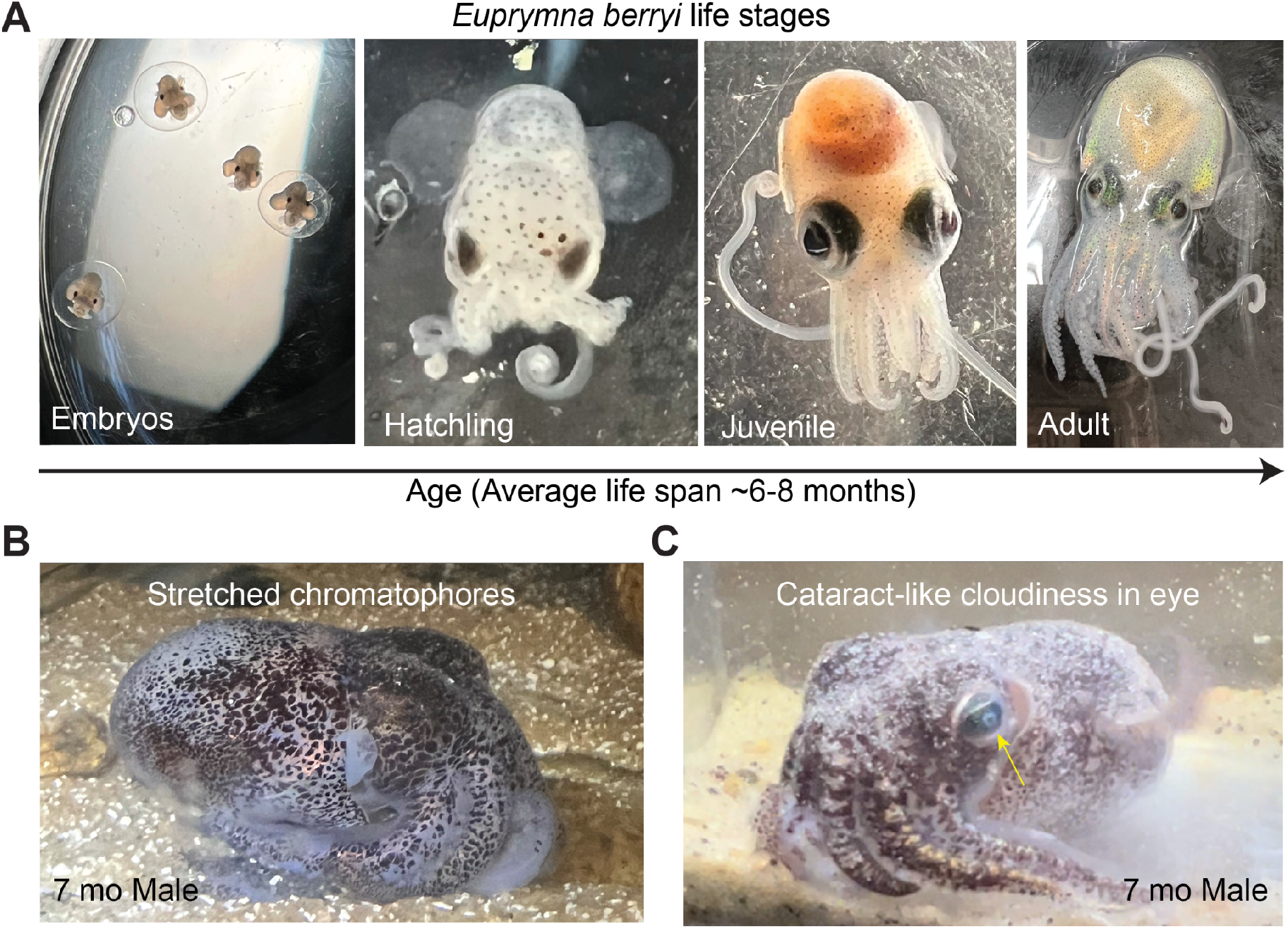
The *Euprymna berryi* squid as an aging model organism. **A**. Four different life stages of *Euprymna berryi*: Embryos, Hatchling, Juvenile, Adult (from left to right). **B**. Stretched chromatophores from a 7-month-old male squid. **C**. Cloudiness in the eye of a senescent 7-month-old male squid (yellow arrow).

The lack of conventional molecular biology tools and experimental methods to study marine invertebrates has proven to be a major roadblock in discovering new biological mechanisms in these organisms. Developing and adapting genetic tools and cell culture systems are necessary to complement genomic studies^8^ in characterizing cell types and will help uncover cellular and molecular mechanisms underlying important functional properties. Despite extensive efforts in establishing compatible marine cell culture systems and techniques to heterologously express exogenous genes, there exists a limited number of examples to date^9,10^. Building upon a cell culture protocol of the Yesso scallop, *Mizuhopecten yessoensis*^9^, we isolate fibroblast-like primary cells from various *E. berryi* tissues of different ages and maintain viable cells in culture. This protocol includes a step-by-step method detailing tissue dissociation, cell isolation, optimized cell culture conditions, and importantly, examples of successful mRNA transfections in isolated cells that can be imaged live or fixed. Because these cells can be isolated from any life stage of the squid, they will allow for future cephalopod and marine aging-related studies^11^ with broader implications of age-dependent changes in cellular and tissue-level organization, physiology, and disease. This work also provides great progress and promise in opening up new strategies for establishing cell culture from a more diverse group of organisms, marine invertebrates and beyond.

## Institutional permissions

All squid husbandry and animal handling, including euthanasia and tissue dissections were performed in accordance with relevant institutional and national guidelines and regulations, i.e. IACUC committees at Princeton University and the Marine Biological Laboratory.

## Key Resources Table

| REAGENT or RESOURCE | SOURCE | IDENTIFIER |
| --- | --- | --- |
| **Antibodies** |  |  |
| Phalloidin-iFluor 488 Reagent (ab176753) | abcam | ab176753 |
| VECTASHIELD® PLUS Antifade Mounting Medium with DAPI | vectashield | H-2000-10 |
| **Bacterial and virus strains** |  |  |
| N/A |  |  |
| **Biological samples** |  |  |
| N/A |  |  |
| **Chemicals, peptides, and recombinant proteins** |  |  |
| Insulin-Transferrin-Selenium (ITS-G) (100x) | Thermo Fisher | 41400045 |
| Leibovitz's L-15 Medium (500 mL) | Thermo Fisher | 11415064 |
| Sodium phosphate monobasic dihydrate | Millipore Sigma | 71500-250G |
| GIBCO Opti-MEM Reduced Serum Medium | Thermo Fisher | 31985062 |
| GIBCO Penicillin-Streptomycin (10,000 U/mL) | Thermo Fisher | 15140122 |
| Fetal Bovine Serum | R&D Systems | S11150H |
| Animal-Free Recombinant Human EGF 100ug | PeproTech | AF-100-15 |
| Recombinant Human FGF-basic (154 a.a.) 100ug | PeproTech | 100-18B |
| Sodium Chloride (Crystalline/Biological, Certified), 500g | Fisher Scientific | S671-500 |
| Trypsin, powder 1:250 | Thermo Fisher Scientific | 27250018 |
| Magnesium sulfate heptahydrate 500g | Sigma Aldrich | M2773-500G |
| Magnesium chloride hexahydrate 500g | Sigma Aldrich | M2670-500G |
| EDTA disodium salt 100g | Sigma | 324503-100G |
| Hoechst 33342 solution | Thermo Fisher | 62249 |
| SPY650-Tubulin | Spirochrome | CY-SC503 |
| Potassium Chloride | Sigma Aldrich | P9541-1KG |
| Sodium Bicarbonate | Sigma Aldrich | S5761-1KG |
| Calcium chloride dihydrate | Sigma Aldrich | C5080-1KG |
| D-(+)-Glucose | Millipore Sigma | G7021-100G |
| **Critical Commercial Assays** |  |  |
| Invitrogen™ Lipofectamine™ MessengerMAX™ Transfection Reagent | Thermo Fisher | LMRNA003 |
| **Experimental models: Organisms/strains** |  |  |
| *Euprymna berryi* squid | Marine Biological Laboratory Cephalopod Program |  |
| **Oligonucleotides** |  |  |
| N/A |  |  |
| **Recombinant DNA** |  |  |
| N/A |  |  |
| **Software and algorithms** |  |  |
| Fiji (ImageJ) | Schindelin et al.<sup>12</sup> | https://imagej.net/software/fiji/downloads |
| **Other** |  |  |
| 24 Well glass bottom plates | Cellvis | P24-1.5H-N |
| Thermo Scientific Nalgene Rapid-Flow Sterile | Thermo Fisher | 0974103 |
| Disposable Filter Units with PES Membrane, 1000mL, Pore Size: 0.20um, |  |  |
| 12x0.20mm Minutens Insect Pins | Indigo Instruments | 33416-20 |
| Insect pins, #3; 38x0.50mm | Indigo Instruments | 33414-3 |
| SYLGARD 184 SILICONE ELASTOMER KIT<br>CLEAR 0.5KG (1.1LB) KIT | Krayden, Inc. | DC4019862 |
| Fine Forceps | Fine Science Tools | 11923-13 |
| Student Fine Scissors, Curved | Fine Science Tools | 91461-12 |

## Materials and equipment setup

For all media and solutions: all steps performed at room temperature (RT) were 21°C, all media and solutions were fixed to osmolality of ∼950 mOsm/kg at a pH of 7.4 and filtered.

Store MBSS at RT. Make DBSS fresh before each dissection or store at 4°C for short-term storage.

Store MCMFS at RT.

Make fresh smaller aliquots (50 ml) of Trypsin solutions A and B right before use and filter:

Store Trypsin-EDTA solutions at 4°C. Warm up to RT before tissue dissociation.

Store Squid growth media A at 4°C. Warm up to RT before use.

Make fresh smaller aliquots (50 ml) of media solutions B-D right before use and filter:

Store Squid growth media B-C at 4°C. Warm up to RT before use.

### Step-by-step method details

To establish squid cell culture, we developed a tissue dissociation and cell isolation protocol building upon a previous study^9^. Our protocol allows isolation of cells from several different tissues, i.e. the optic lobes, eyes, and gills (Figure 2A-C) that provide the most successful cultures (Figure 2D-F). We can also isolate cells from the skin and arm, but not as robustly as the aforementioned tissues.

**Figure 2:**
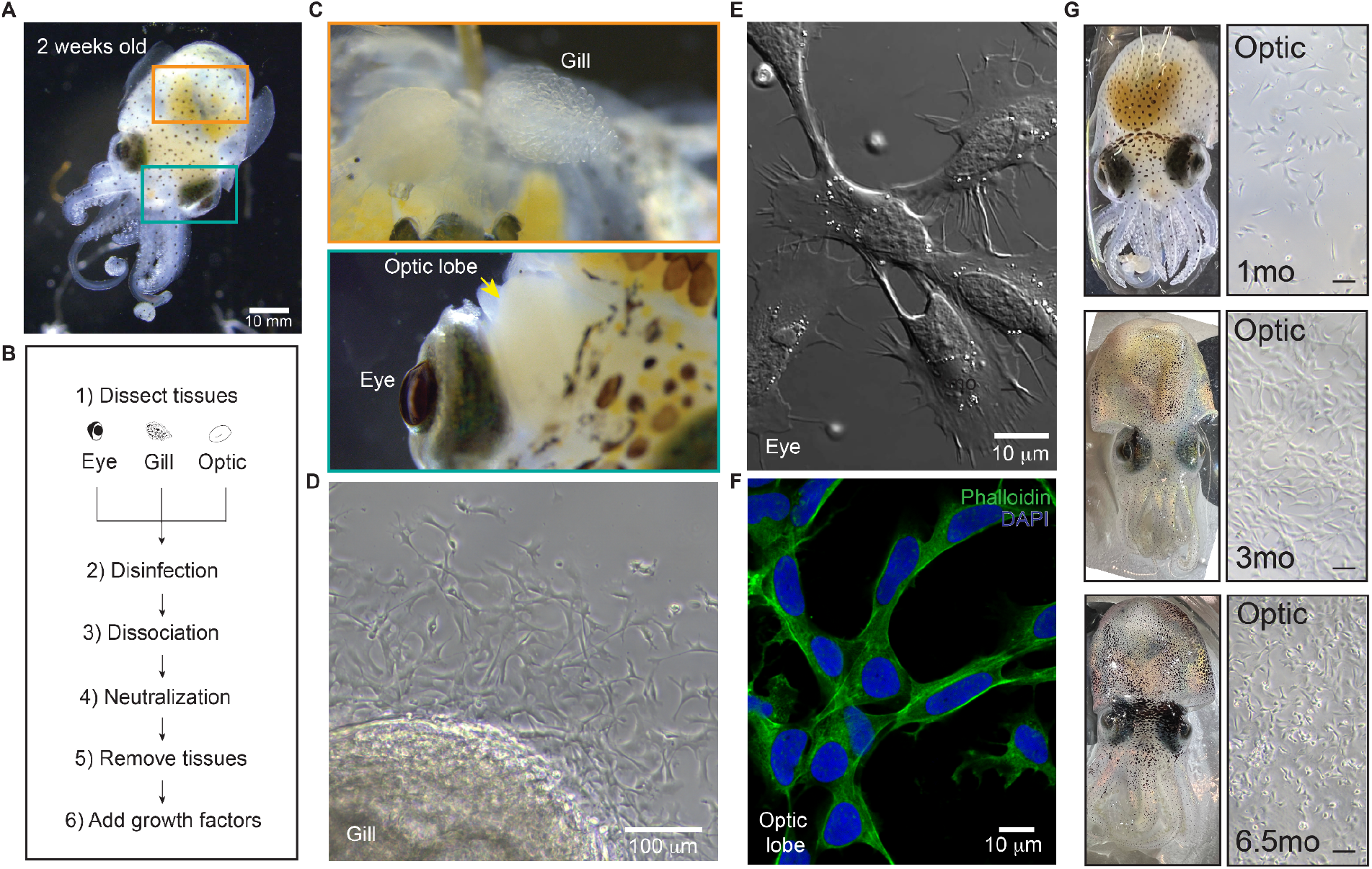
Squid cells isolated from various tissues and life stages. **A**. 2-weeks-old *E. berryi* hatchling. **B**. Workflow for primary cell isolation from eyes, gills, and optic lobes from any life stage. **C**. Close-up image of the gill found in the yellow boxed region and the eye and optic lobe (yellow arrow) in the turquoise boxed region of the squid in A. **D**. Images of cells migrating out from an *E. berryi* gill. **E**. DIC image of cells isolated from an *E. berryi* eye. **F**. Optic lobe cells were fixed and stained with phalloidin (green) and DAPI (blue) to mark the nuclei. **G**. Optic lobe cells isolated from 1-month-old, 3-month-old, 6.5-month-old squids (squids not shown to scale). All three scale bars indicate 50 µm.

### Primary cell isolation and culturing conditions

Timing: 2-3 hours

### Materials and preparations before starting

#### Dissecting tools

- Sylgard plate
- Insect pins
- Autoclave dissecting tools, i.e. forceps, scissors

*Media* (see ‘Materials and equipment setup’ section for recipes)

Before the dissection

1. Make the Disinfection Balanced Salt Solution (DBSS) from^9^ (Table 1) by adding 0.5% penicillin-streptomycin to the Molluscan Balanced Salt Solution (MBSS) from^9^ (Table 1). All cells are plated in 24 well plates unless noted. Add enough volume of DBSS to fully cover each piece of tissue (500 ul into each well of a 24 well plate that will contain optic lobes or gills. Add 1 ml to wells that will contain eyes). Make sure all media and solutions are at RT before dissecting the squid and subsequently plating cells isolated from the squid.
2. Autoclave all dissection tools.
3. Spray down all equipment, bench area, and dissection tools with 70% ethanol. Use good sterile aseptic technique.

**Table 1:** Molluscan Balanced Salt Solution (MBSS) from^9^.

| Table 1: Molluscan Balanced Salt Solution (MBSS) from<sup>9</sup> |  |
| --- | --- |
| NaCl | 26.0 g |
| KCl | 1.08 g |
| MgCl<sub>2</sub>6H<sub>2</sub>O | 4.70 g |
| MgSO<sub>4</sub>7H<sub>2</sub>O | 6.52 g |
| NaHCO<sub>3</sub> | 0.30 g |
| NaH<sub>2</sub>PO<sub>4</sub>2H<sub>2</sub>O | 0.06 g |
| CaCl<sub>2</sub>2H<sub>2</sub>O | 1.48 g |
| Glucose | 0.3 g |
| Dissolve in 1 L dH<sub>2</sub>O |  |
| Add 0.5% Penicillin-streptomycin and filter to make **Disinfection Balanced Salt Solution (DBSS) from<sup>9</sup>** (for tissue disinfection) |  |

Squid dissection

1. Perform euthanasia on squids by slowly adding ethanol into tank seawater^4,13^. In small increments, slowly add in more ethanol up to 5% total maximum for up to 30 minutes^4,13^.
2. Once the animal has stopped respiring, remove the animal from the ethanol solution and place on a sylgard plate.
3. Use two pins to secure the squid on the sylgard by pinning down the top of the mantle and another pin down near the beak (Figure 3A).
4. Use one forcep and a scissor to make an incision around the back of the eye and cut enough of the skin to remove the eye. Place the eye in a well in the 24 well plate containing DBSS.
5. Next, remove the whole optic lobe (behind the eye) along with pieces of connective tissue (Figure 3B) (this is critical as the connective tissue seems to hold a lot of cells) and place them into the 24 well plate containing DBSS.
6. Repeat for the other side of the animal to dissect the remaining eye and optic lobe.
7. To remove the gills, turn the animal over so that its back is facing up and make a clean scission right under the bottom of the mantle/above the central mass.
8. Make an incision down the ventral mantle and open and pin down the flaps (Figure 3C).
9. Remove each gill and wash the tissue in the 24 well plate in DBSS.

**Figure 3:**
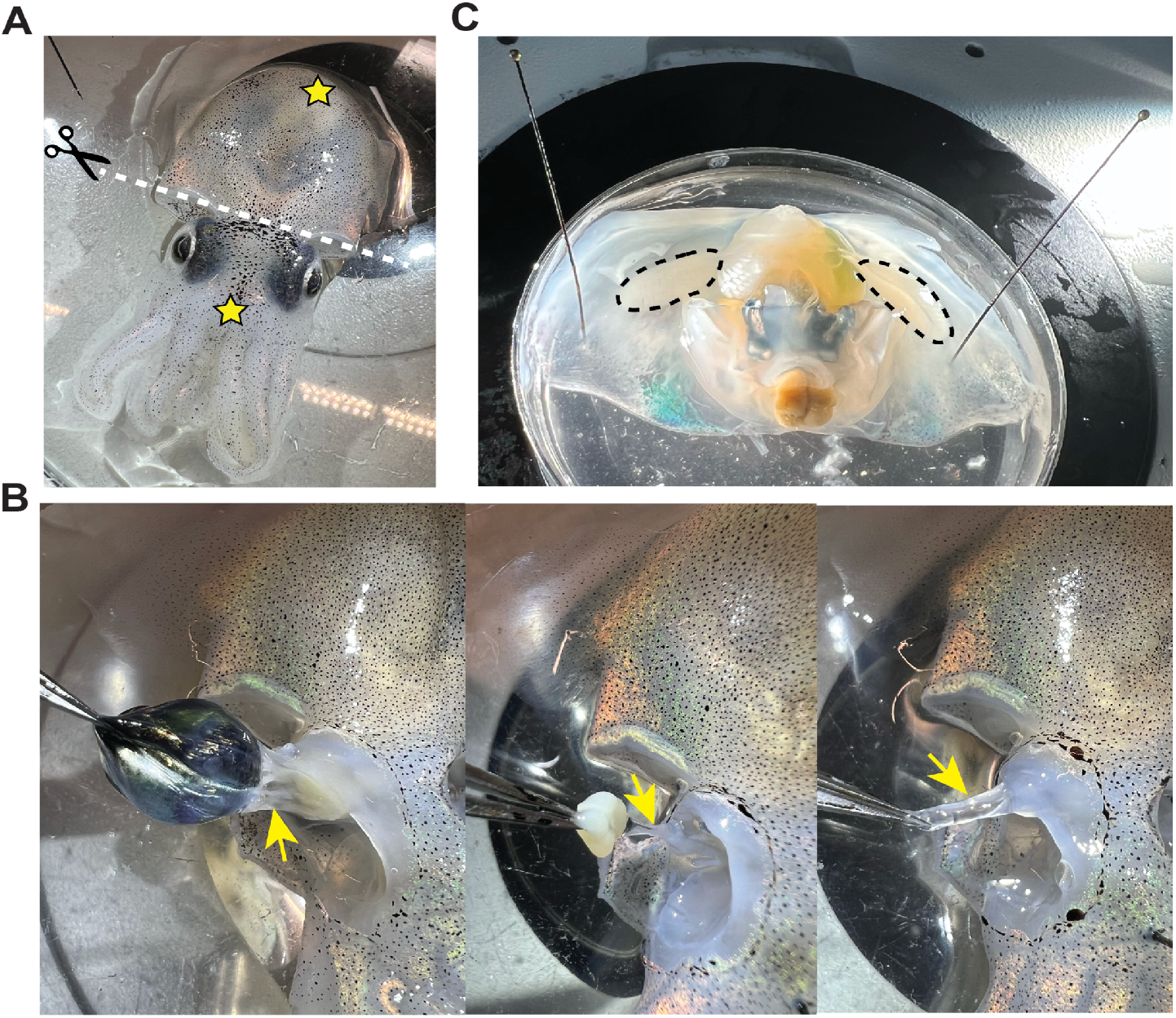
Squid dissection up close. **A**. Euthanized adult squid on a sylgard plate. Yellow stars indicate where to pin down the squid with two insect pins. White dotted line shows where to make the scission to separate the mantle off from the rest of the body. **B**. When dissecting out the eye (left) and optic lobe (center), make sure to also remove as much of the connective tissue mass (right and in yellow arrows) and add to the well plate. **C**. Once opening up the mantle, the gills are to the left and right as labeled by the black dotted lines.

Tissue dissociation and cell isolation:

1. Leave the tissues in DBSS for 30 minutes to disinfect. Gently shake or rock the plate in 5-10 minute intervals.
2. Remove the tissues from the DBSS solution and move into another clean well with a sterile forcep. Leave the DBSS solution in the well as there are cells dissociated already from the disinfection step.
3. To dissociate cells from the tissue, add enough trypsin-EDTA solution A (Table 3; made from the Molluscan Calcium/Magnesium Free Solution (MCMFS) from^9^, Table 2) to fully submerge the tissue for 10 minutes (500 ul for optic lobe and gill, 1 ml for an adult eye).
4. Neutralize the trypsin-EDTA by adding 2 ml of Squid Growth Media B (Table 5; made from Squid Growth Media A stock, Table 4) and leave with the tissue for 15 minutes.
5. Remove the tissue with clean forceps and place into a new well with Squid Growth Media C (Table 5) for 1 hour.
6. Remove the tissue with clean forceps and place into a new well with Squid Growth Media D (Table 5) overnight.
7. All wells from this section can be used for downstream applications - important to wash out the DBSS, trypsin-EDTA wells with Squid Growth Media C or MCMFS (Table 2) and replenish with Squid Growth Media D (daily).
8. Cells are left at RT and readily adhere to plates even as early as ∼1-2 hours post cell isolation and are viable for 3-5 days. Once fully attached to the plate, they are ready for use! In a given 24-well plate, the number of cells isolated in a single well usually range from 0.5 - 2 × 10^5^ cells.
9. Cells can be lifted by manually scraping with cell scrapers or using the trypsin-EDTA solution B (Tables 3) and plated in other dishes (Figure 4A) or collected for further downstream application, such as DNA and RNA isolation.

**Table 2:** Molluscan Calcium/Magnesium Free Solution (MCMFS) from^9^.

Store MBSS at RT. Make DBSS fresh before each dissection or store at 4°C for short-term storage.
| **Table 2: Molluscan Calcium/Magnesium Free Solution (MCMFS) from<sup>9</sup>** |  |
| --- | --- |
| NaCl | 26.0 g |
| KCl | 1.08 g |
| NaHCO<sub>3</sub> | 0.30 g |
| NaH<sub>2</sub>PO<sub>4</sub>2H<sub>2</sub>O | 0.06 g |
| Glucose | 0.3 g |
| Dissolve in 1 L dH<sub>2</sub>O |  |

**Table 3:** Trypsin-EDTA for tissue dissociation and passaging cells.

Store MCMFS at RT. Make fresh smaller aliquots (50 ml) of Trypsin solutions A and B right before use and filter:
| **Table 3: Trypsin-EDTA for tissue dissociation and passaging cells** |  |
| --- | --- |
| **Trypsin-EDTA solution A** (for tissue dissociation): | Add 0.25% Trypsin powder and 0.02% EDTA powder to MCMFS |
| **Trypsin-EDTA solution B** (for lifting cells off plates and dishes): | Add 0.05% Trypsin powder and 0.02% EDTA powder to MCMFS |

**Table 4:** Squid growth media A from^9^ (use as stock; no antibiotic and no serum)

Store Trypsin-EDTA solutions at 4°C. Warm up to RT before tissue dissociation.
|  |  |
| --- | --- |
| NaCl | 19.67 g |
| KCl | 0.81 g |
| MgCl<sub>2</sub>6H<sub>2</sub>O | 3.53 g |
| MgSO<sub>4</sub>7H<sub>2</sub>O | 4.89 g |
| NaHCO<sub>3</sub> | 0.23 g |
| NaH<sub>2</sub>PO<sub>4</sub>2H<sub>2</sub>O | 0.044 g |
| CaCl<sub>2</sub>2H<sub>2</sub>O | 1.11 g |
| Glucose | 0.23 g |
| Dissolve in 1 L of Leibovitz's media (L-15) |  |

**Table 5:** Squid growth media B-D.

Store Squid growth media A at 4°C. Warm up to RT before use. Make fresh smaller aliquots (50 ml) of media solutions B-D right before use and filter:
| **Table 5: Squid growth media B-D** |  |
| --- | --- |
| **Squid growth media B**<br>(antibiotic and serum to neutralize trypsin) | 0.5% Penicillin-streptomycin<br>8% L-15 media<br>10% Fetal Bovine Serum<br>81.5% Squid growth media A |
| **Squid growth media C**<br>(antibiotic only and no serum) | 0.5% Penicillin-streptomycin<br>8% L-15 media<br>91.5% Squid growth media A |
| **Squid growth media D**<br>(antibiotic and growth factors) | 0.5% Penicillin-streptomycin<br>8% L-15 media<br>1% Insulin-Transferrin-Selenium (ITS-G)<br>10 ng/ml of the fibroblast growth factor (FGF)<br>100 ng/ml of the epidermal growth factor (EGF)<br>90.5% Squid growth media A |

**Figure 4:**
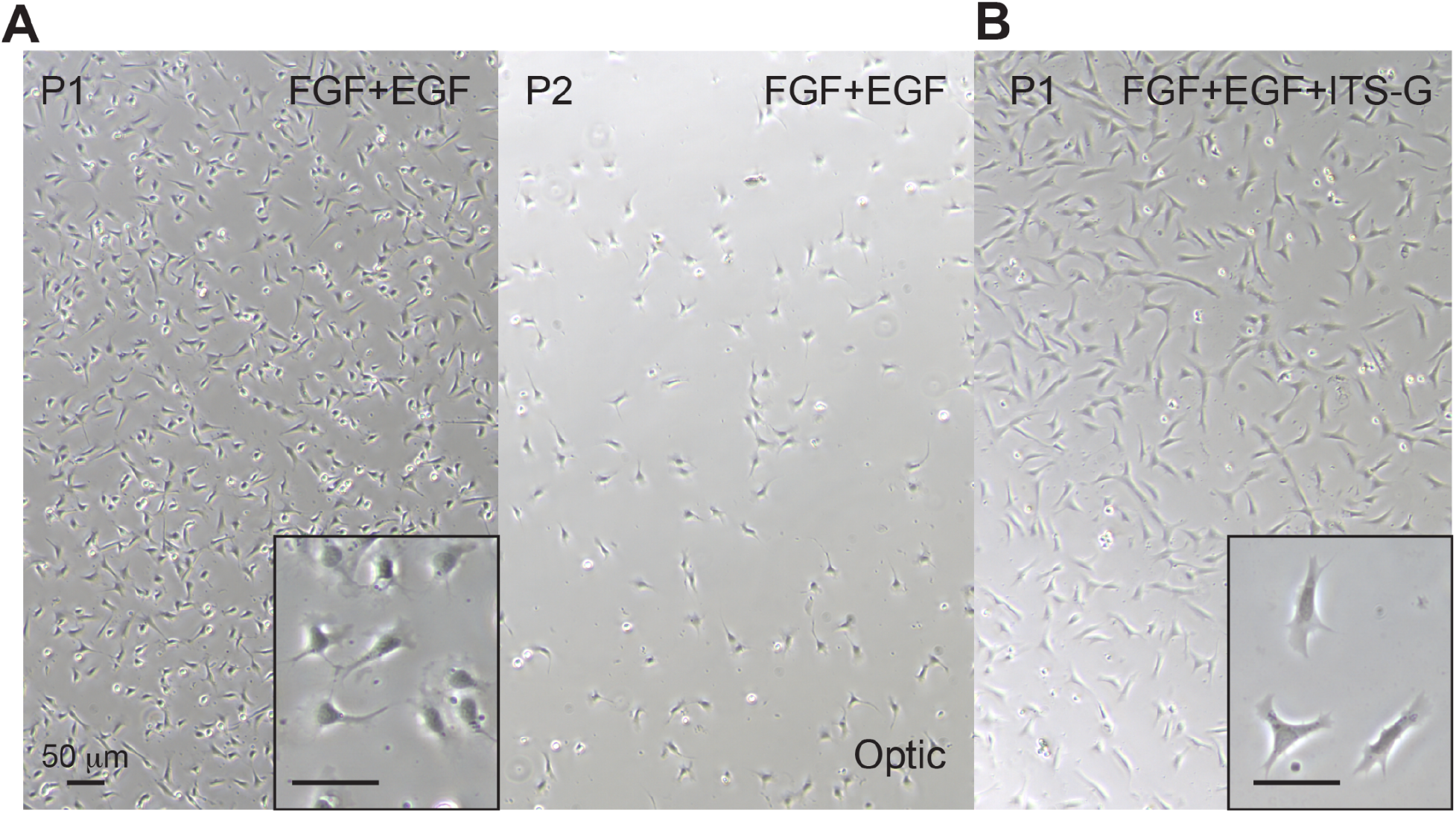
Improving squid cell culture conditions and broader utility. **A**. Squid optic lobe cells can be scraped and replated (P0 to P1 to P2), with cells readily re-adhering to a glass bottom dish, although there is loss in cell number (P2). **B**. Adding ITS-G to squid growth media with FGF+EGF improves cell attachment. Insets show close-ups of cells from the same well for each condition shown. Scale bars in insets indicate 50 µm.

## mRNA transfections and live cell imaging

Timing: Following Lipofectamine MessengerMAX manufacturer’s protocol, Preparation/Incubation: 20 minutes

Final incubation: 1 day

1. Plate cells to be 70-90% confluent at the time of transfection.
2. Wash cells twice with Squid Growth Media C but leave out the penicillin-streptomycin. Leave the cells in this growth media prior to transfection.
3. Prepare mRNA (500 ng)-lipid complexes following Lipofectamine MessengerMAX manufacturer’s protocol. Complexes made in ‘no serum Opti-MEM’ (osmolality doesn’t have to be fixed).
4. Add complexes to cells left in Squid Growth Media C with no penicillin-streptomycin for 24 hours at RT and check expression on a microscope.
5. Hoechst can be added to live cells to also mark the nuclei. Use a 1:50,000 Hoechst dilution in growth media and leave to incorporate for 1 hour before imaging. Remove media with Hoechst, wash two times and replenish with fresh growth media. All live imaging should be conducted at RT with ambient-level CO_2_.

## Cell fixation

Preparation before starting:

Make fresh 4% PFA in MBSS at RT

Timing: 30 minutes

1. Remove growth media and wash cells with MBSS twice to wash out the growth media.
2. Add 4% PFA in MBSS and incubate at RT for 10-15 minutes.
3. Perform three sequential 5 min washes with MBSS.
4. Leave cells in MBSS at RT if using immediately or store in 4°C.
5. Perform Immunofluorescence/FISH using permeabilization buffer with 0.5% Triton-X. Add 1:1000 Hoechst dilution in MBSS for 15 minutes. Wash out with MBSS.

## Expected outcomes

After modifying media conditions from^9^ by optimizing osmolality levels and adding ITS and growth factors FGF/EGF, fibroblast-like cells from the outer layers of the tissue adhered to the bottom of a glass bottom dish (Figure 2D). These cells take on fibroblast-like characteristics and morphologies, consistent with reports from a previous study that characterized a population of fibroblast-like cells from the outer layer of the optic lobes from whole tissue RNA-seq^14^.

Additionally, these isolated cells are adherent and exhibit elongated shapes with flattened nuclei and cytoplasmic projections (Figure 2E, 5A-B). Nuclei of cells exhibit a variety of morphologies (Figure 5B), and tubulin stainings show spindly microtubules and an interesting unknown cytoplasmic body adjacent to nuclei (Figure 5A). While cells would readily crawl out of explants after being plated in these optimized media conditions (Figure 6A-B), cells were maintained for longer *in vitro* once the explant was removed as small remnants of tissue can harbor small microorganisms and contaminants (Figure 6B) that lead to cell death even with the presence of antibiotics like penicillin-streptomycin, as also seen in previous studies^9^. Cell growth and proliferation rates were difficult to assess even after removing explants due to smaller fragments of tissues remaining in culture^9^. Instead of plating the explants on the dish, leaving the explants afloat led to more prolific and robust cell isolation and easier removal of tissue and performing subsequent washes.

**Figure 5:**
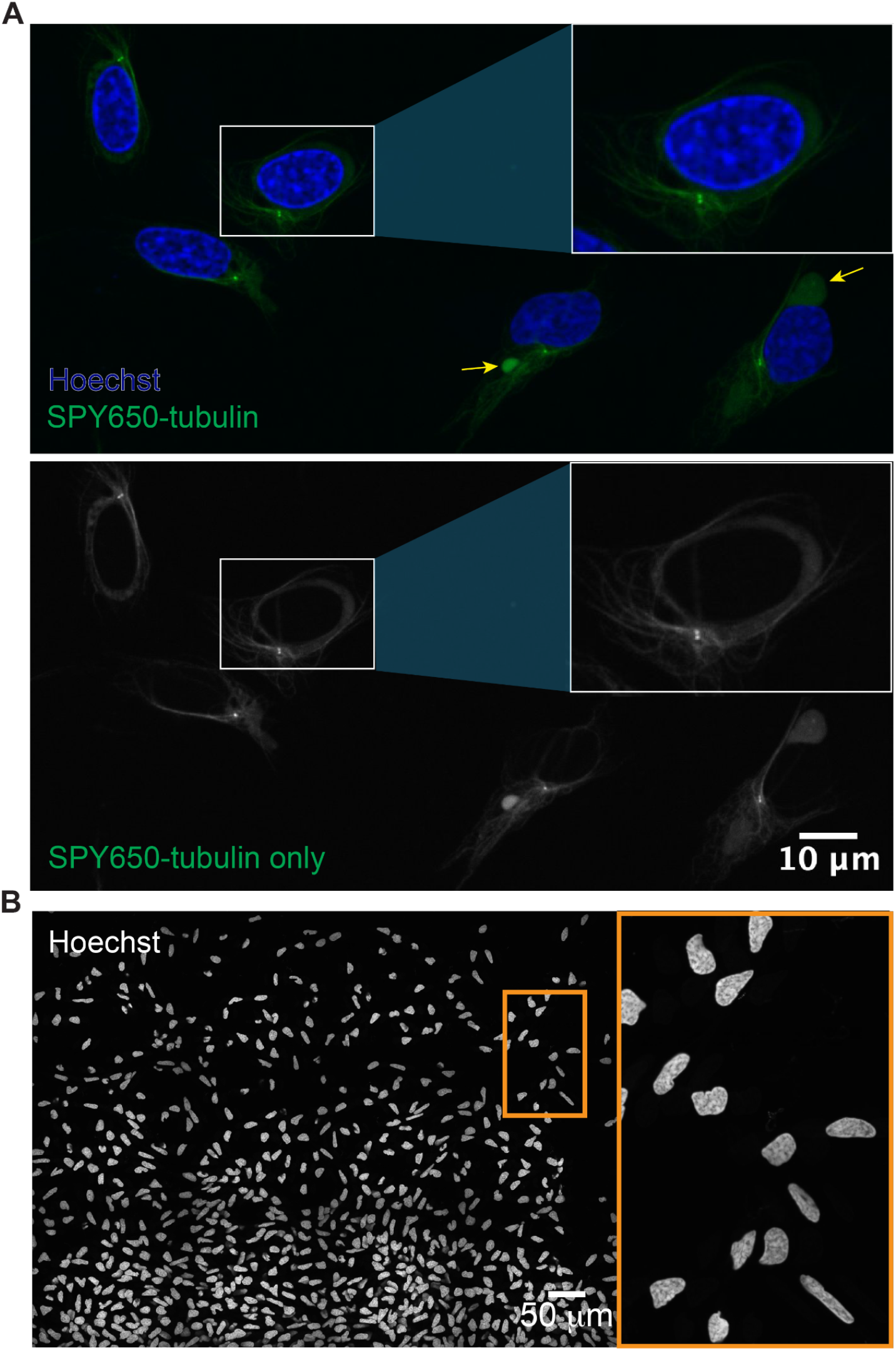
Squid cells are fibroblast-like. **A**. Optic lobe cells labeled with Hoechst (blue) to stain for nuclei and SPY650-tubulin (green) to label microtubules live. Yellow arrows indicate cytoplasmic bodies. Insets show close-ups of the cell in the white box. **B**. Large image of Hoechst-labeled squid nuclei with the orange box showing heterogeneity of nuclear shape.

**Figure 6:**
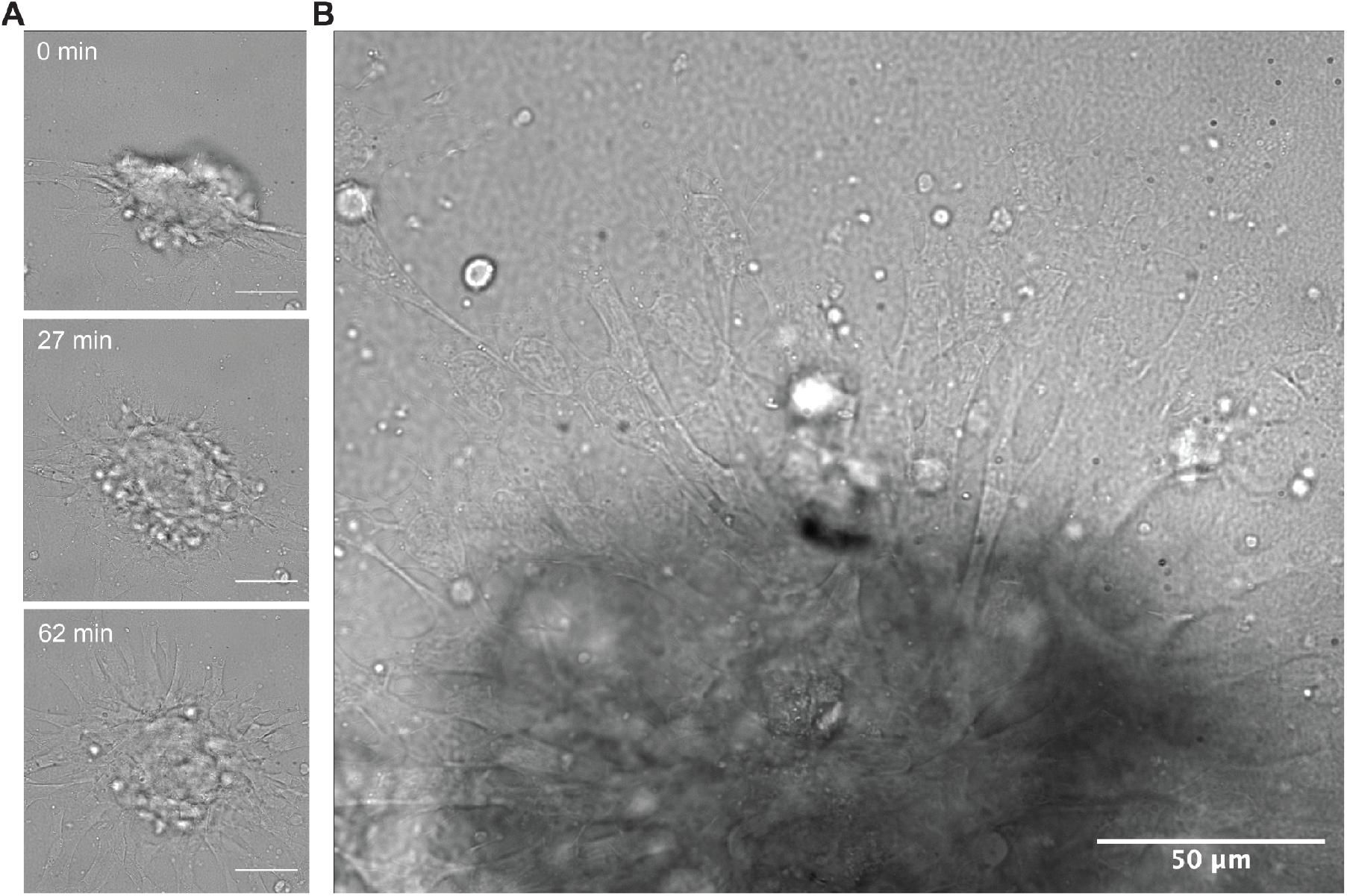
Cells migrate out from tissue explants. **A**. Small piece of squid tissue explant from an optic lobe adhered on glass bottom, with panels showing cells crawling out of an explant over the course of roughly an hour (62 minutes). Scale bar indicates 50µm. **B**. Cells readily migrate out from tissue explant along with microorganisms from the tissue.

### mRNA transfections

We next show successful transfection of exogenous genes into optic lobe cells (Figure 7). Using lipofection-based standard methods commonly used for mammalian cell culture, we were able to express mRNAs in squid cells. Only a small subset of cells expressed NPM1-RFP of *Xenopus laevis* sequence (Figure 7A-B). FIB1-eGFP of *Xenopus* sequence and Kaede NLS-GFP also expressed in squid nuclei (Figure 7C). Both NPM1 and FIB1 formed 1-2 bright puncta towards the peripheral areas in the nucleus (Figure 7B-C), different from most nucleolar morphologies. However, these puncta resemble the NORs that were detected using silver nitrate (Ag)-staining in interphase octopus (*C*.*chinensis*) cells^15^. 24 hours post transfection, NPM1-RFP mRNA expressed 5.17% of 329 nuclei, FIB1-eGFP expressed in 3.70% of 379 nuclei, and Kaede NLS-GFP in 1.21% of 1070 nuclei (Figure 7E). Future work will confirm whether these puncta are indeed squid nucleoli and will also investigate whether they form a multilayered structure in squids as they do in mammalian cells^16^. Successfully expressing exogenous genes will further help characterize nuclear and cytoplasmic compartments in squid cells.

**Figure 7:**
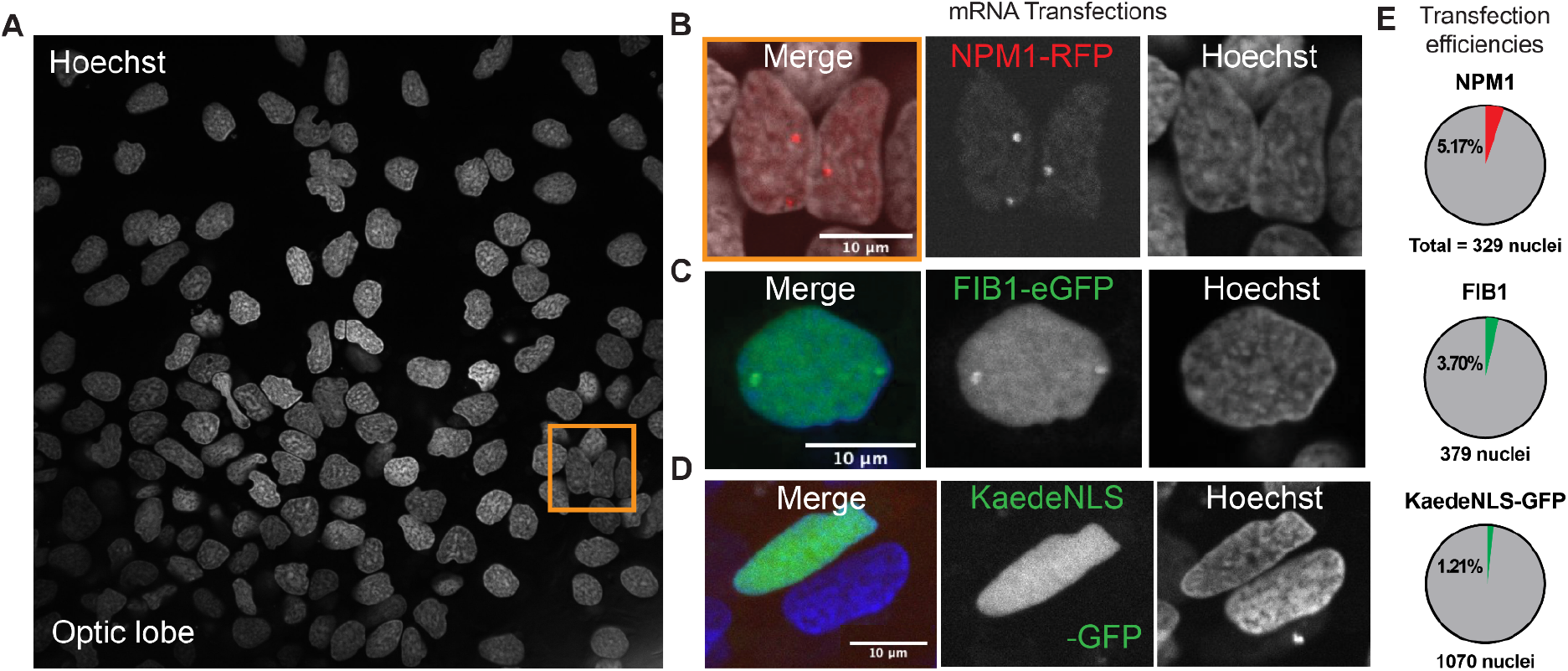
Squid cells express exogenous genes. **A**. Squid optic lobe cells (cells isolated and plated 1 day prior to transfection) labeled with Hoechst **B**. express NPM1-RFP of *Xenopus laevis* sequence (orange box in A), **C**. FIB1-eGFP also of *Xenopus laevis* sequence, and **D**. KaedeNLS-GFP 24 hours post mRNA transfection. **E**. Transfection efficiencies of mRNAs transfected in B-D

## Limitations

While mRNA transfections were successful, DNA plasmids were not able to be expressed in these cells despite having tried various methods, including with the use of lipofection-based reagents, electroporation, lentiviral vectors, AAV vectors, along with various promoters, i.e. CMV, Ubc, SFFV, and OsHV among others (data not shown). One reason why it is difficult to express DNA plasmids in these cells could be that the promoters used here are not compatible with squid cells. Also, squid cells could have a robust line of defense against foreign DNA. Another reason could be due to low proliferation rates, i.e. we have not been able to observe a single cell division event, since methods like electroporation and lipofection require cell division for the DNA to enter the nucleus. Thus, microinjection or nucleofection may serve as promising alternatives. These methods can allow for transfection of non-dividing cells. Most reagents used in this study have been optimized for mammalian cell investigations, including the supplemental growth factors, thus there is a need for tool development to utilize antibodies, growth factors and promoters optimized for molluscan cells. Future studies will address these points.

In summary, we have established primary cell culture from *Euprymna berryi* squid tissues. Using an optimized protocol for tissue dissociation and cell isolation, fibroblast-like adherent cells from various tissues of different age groups migrate out from explants and form monolayer of cells *in vitro*. Future studies will employ single-cell RNA sequencing to characterize the cell types isolated from the tissue along with cells that are maintained in culture with the specific culture conditions described here. This work allows for a simplified and controlled source of cells for investigations into marine and cell biology, along with future progress in establishing a squid cell line could help reduce the number of animals sacrificed for experimentation^17^.

## Troubleshooting

### Problem 1

There is excessive debris from the tissue and/or contaminants and microbes in culture.

### Potential solution

Instead of making an incision shown in Figure 3A to separate off the mantle from the rest of the body, proceed by dissecting out relevant tissues of an intact squid and remove tissues. Add additional wash steps using Squid growth media C or D.

### Problem 2

Not enough cells dissociating from tissues.

### Potential solution

Gently pipette up and down the Trypsin-EDTA solution A during the tissue dissociation step and/or the Squid growth media B during the trypsin neutralization step (make sure to do this very gently or else excess tissue will dissociate).

### Problem 3

Hoechst/DNA dyes do not incorporate into cells.

### Potential solution

Try a lower dilution and leave in wells for longer.

### Problem 4

mRNAs are not expressing in squid cells.

### Potential solution

Leave in the mRNA-lipid complexes for another 24 hours. If the cells start looking unhealthy, try washing out the complex+media with fresh Squid Growth Media C with no penicillin-streptomycin to let cells recover. Another option is to try another round of transfections by adding newly made mRNA-lipid complexes after the first 24-hour mark.

## Resource availability

### Lead contact

Further information and requests for any resources and reagents should be directed to and will be fulfilled by the lead contact, Cliff Brangwynne.

## Technical contact

Questions regarding technical specifics of the protocol should be directed to the technical contact, Yoonji Kim.

## Materials availability

This study did not generate new unique reagents.

## Data and code availability

This study did not generate code. Raw data reported in this paper will be shared by the lead contact upon request.

## Acknowledgements

We thank the Marine Biological Laboratory’s Cephalopod Program, in particular Bret Grasse, Taylor Sakmar, and Miranda Vogt for all assistance and guidance on animal husbandry training and advice, euthanasia protocol, and supplying squid; the Princeton Laboratory Animal Resources department, in particular Lauren Shouey, Katherine Wu, Hannah Westin, Phil Johnson for animal husbandry related support; the IACUC committees both at Princeton University and the Marine Biological Laboratory, Carrie Albertin and Gjenni Voss for their helpful advice; Deepa Rajan and Kirby Leo for performing preliminary experiments and imaging during the MBL Physiology course; Holly Cheng for the NPM1-RFP/FIB1-eGFP mRNAs and for acquiring/denoising Figure 2E; the Rosenthal lab for the Kaede-NLS-GFP mRNA; and Jing Xia for sharing reagents and providing help with the osmometer and performing calibrations. We also thank Aalok Varma, Loren Looger, Helen Farrants, Aaron Lin, and Sofi Quinodoz for helpful discussions and providing reagents for troubleshooting DNA plasmid transfection experiments; Nick Treen for bringing the Yoon et al. 2022 paper to our attention; Andrea Bodnar for helpful discussions regarding cell proliferation; Yan Wang for insights on cephalopod aging; Evangelos Gatzogiannis for microscopy assistance, and the rest of the Brangwynne lab for helpful discussions.

This work was supported by the Marine Biological Laboratory’s Physiology course, the Marine Biological Laboratory’s Whitman Center Fellowship, and the Howard Hughes Medical Institute (C.P.B.). Y.K. was supported by the NSF GRFP (DGE-2039656). J.J.C.R. is funded by the RoL:NSF-BSF:IMAGiNE (2110074) and the NSF EDGE FGT (2220587).

## Author Contributions

Y.K. and C.P.B. designed the study. Y.K. optimized squid primary cell culture conditions. Y.K. and H.M.T. performed squid dissections, experiments and analysis, with advice from J.J.C.R. and C.P.B. J.J.C.R. trained Y.K. in squid dissections, and Y.K. trained H.M.T. Y.K. and the Laboratory Animal Resources at Princeton University managed squids of all life stages. Y.K. and C.P.B. wrote the manuscript, and Y.K. made the figures, with contributions from all authors.

## Additional Materials and Methods

### Squid Images and Dissections

A Leica fully motorized M205FA stereomicroscope was used for squid hatchling dissections. This stereomicroscope was equipped with a Plan Apo 1.0x air lens and a Leica Arc Lamp white light source for fluorescence with GFP and mCherry filter cubes. Images were recorded with a Leica FlexaCam C3 CMOS camera. Images of all squids in Figure 1, juvenile and adult animals before dissection in Figure 2G, and close-ups of squid dissections in Figure 3 were taken with an iPhone 13 Pro.

### Microscopy and Live Cell Imaging

A Nikon spinning disk confocal microscope, consisting of a Yokogawa W1 SoRa spinning disk confocal scanhead equipped with the Yokogawa W1 bypass mode, was used for fluorescence and DIC microscopy. This system was built around a Nikon Ti2-E fully motorized microscope harboring dual Hamamatsu Fusion BT sCMOS cameras. A Nikon CFI Plan Apo Lambda D 60x oil (MRD71670) was used with 405, 488, 561, 640 nm lasers and transmission light. We used a Mad City Labs piezo z stage for acquiring z-stack images. Some images were acquired using a Nikon A1 laser scanning confocal microscope with a 60X oil-immersion objective (1.4 numerical aperture) using DIC and were denoised using Denoise.ai.

### mRNA transfections

NPM1-RFP and FIB1-eGFP mRNA’s were gifts from Holly Cheng. The Kaede NLS-GFP mRNA was a gift from the Rosenthal Lab. Lipofectamine MessengerMAX transfection reagent (Thermo Fisher, LMRNA003), following the manufacturer’s instructions, was used to transfect squid cells using these mRNAs. Cells were imaged 24 hrs post transfection.

### Measuring osmolality

A vapor pressure osmometer (Knauer K-7000) was used to measure osmolality of the squid culture media optimized in this study, using manufacturer’s guidelines to perform measurements and calibrations. Briefly, the osmometer was calibrated using a standard 100, 300, 500, 900, 1100 mOsmol/kg NaCl set of solutions. Pure solvent droplets were first applied to both thermistors to set the baseline. The “Autozero” function was then applied. A droplet of each standard solution was then applied to the sample thermistor, and the bridge voltage was adjusted until the displayed measurement matched the known osmolality of the standard. Calibration was repeated three times to ensure consistency. Once calibrated, droplets of the culture media and buffers were then applied to measure osmolality.

